# The Readiness Potential reflects the internal source of action, rather than decision uncertainty

**DOI:** 10.1101/782813

**Authors:** Eoin Travers, Patrick Haggard

**Affiliations:** Institute of Cognitive Neuroscience, University College London

**Keywords:** Voluntary action, Readiness Potential, Decision making

## Abstract

Voluntary actions are preceded by a Readiness Potential (RP), a slow EEG component generated in medial frontal cortical areas. The RP is classically thought to be specific to internally-driven decisions to act, and to reflect post-decision motor preparation. Recent work suggests instead that it may reflect noise or conflict during the decision itself, with internally-driven decisions tending to be more random, more conflictual and thus more uncertain than externally-driven actions. To contrast accounts based on endogenicity with accounts based on uncertainty, we recorded EEG in a task where participants decided to act or withhold action to accept or reject visually-presented gambles, and used multivariate methods to extract an RP-like component.. We found no difference in amplitude of this component between actions driven by strong versus weak evidence, suggesting that the RP may not reflect uncertainty. In contrast, the same RP-like component showed higher amplitudes prior to actions performed without any external evidence (*guesses*) than for actions performed in response to equivocal, conflicting evidence. This supports the view that the RP reflects the internal source of action, rather than decision uncertainty.

**Significance Statement:** The EEG Readiness Potential (RP) seen prior to self-initiated actions is often taken as a neural marker of volition (Shibasaki & Hallett, 2006). It has been argued that the RP may instead reflect conflict and uncertainty in the decisions leading up to these actions, raising questions about decades of previous findings. We directly tested these explanations by asking participants to decide whether to act or not in a gambling task, while manipulating both uncertainty and the source of the information (internal or external) guiding the decision. Only the source of information affected the presence of the RP. Therefore, the RP remains a marker of internally-generated voluntary action, not a marker of uncertainty.

## Introduction

According to one influential definition, actions are voluntary if they are produced not in response to an external trigger, but from some internal source (Passingham et al., 2010). Such voluntary actions are often preceded by the *Readiness Potential* (RP): a slow negative EEG component recorded over midline electrodes in the second or so prior to action, generated primarily by the Supplementary Motor Area (SMA; Kornhuber & Deecke, 1965; Shibasaki & Hallett, 2006). The RP has often been considered a marker of the internal source of voluntary action, or at least a consequence of the operation of that source. An average RP deflection can be seen prior to the time at which participants report being aware of a decision to move, raising questions about the role of conscious intentions in action (Libet, 1985). However, it is not yet clear what the RP reflects.

One view, based on neuroanatomical and neurophysiological studies of the frontal cortex, is that there are two distinct neural pathways for action. The lateral pathway, for exogenous stimulus-driven actions, connects sensory regions to the primary motor cortex (M1) via the lateral premotor cortex. The medial pathway, for internally-generated endogenous actions, connects the prefrontal cortex to M1 via the medially-located SMA (Passingham, 1993; Passingham et al., 2010). This proposal is supported by functional neuroimaging results comparing simple self-initiated movements to externally-triggered movements, or to baseline (Frith et al., 1991; Jahanshahi et al., 1995), and source localisation of the RP (Shibasaki & Hallett, 2006). There is also evidence from non-human primates that lesions to medial premotor pathways impair self-initiated actions while leaving responses to stimuli intact (Passingham, 1993; Thaler et al., 1995). However, direct recordings from rodent motor regions suggest self-initiated actions require the same computations as externally-triggered actions, but at a slower timescale (Elsayed et al., 2016; Lara et al., 2018).

An alternative view is that the RP reflects uncertainty in the decision that leads to actions. Nachev and colleagues (Nachev et al., 2005, 2008) propose that SMA and pre-SMA activation occurs when there is conflict or insufficient constraints in decisions about when to act. This is in contrast to the neighbouring anterior cingulate cortex, which is activated when there *is* conflict in decisions about what action to perform (Botvinick et al., 2004). Nachev and colleagues suggest that the SMA activity (and hence the RP) during voluntary action tasks is not due to the self-initiated nature of movement, but rather due to the fact that the decision to move is under-determinedby external evidence, leading to competition and conflict (Botvinick, Cohen, & Carter, 2004). Thus, the current literature suggests at least two possible explanations of the RP: the RP either reflects that an action is produced endogenously, or it reflects uncertainty and noise in the decision leading to action.

Schurger, Sitt and Dehaene (2012) showed that the RP could be produced by a noisy evidence accumulation process when the input signal is very weak. In this case, random noise in the process eventually passes the threshold for action. When multiple instances of accumulated random noise are averaged together, time locked to when the threshold is crossed, this model reproduces the slow ramping shape of the RP. Schurger et al. (2012) do not consider whether this is the same evidence accumulation process as is involved in decisions based on external cues. On one view, the same mechanism might produce both self-initiated actions and externally-triggered decisions (O’Connell et al., 2018), but with a very weak input signal for self-initiated actions. Since the RP is reduced or absent prior to externally-trigged actions – actions with strong inputs – compared to self-initiated actions (Jahanshahi et al., 1995) – actions with weak inputs – it would follow that the RP reflects this weakness in the input. For example, RP amplitudes should be inversely proportional to the strength of sensory evidence for action. This interpretation would be consistent with what is proposed by Nachev and colleagues. Alternatively, there may be two separate generators for action (Passingham, 1993), and this noise accumulation model only describes the mechanism for internally-generated actions (Maoz et al., 2019). In this case, the RP should be stronger for internally-generated actions, and should be unaffected by the strength of any external sensory signals to act. This would be consistent with the classic view that the RP reflects the endogenicity of action.

To test these competing theories, we designed a gambling task that varies both whether participants’ actions are driven by exogenous evidence or endogenous cues, and also how uncertain participants are about whether or not to act. On each trial, participants were visually presented with a gamble, and decided to either act or refrain from acting in order to accept or decline it.

On most trials, a specific combination of gamble outcome value and success probability was shown, so participants could this exogenous information to decide whether or not to act. However, on some trials this information was hidden from participants, so participants had to guess, relying on endogenous cues, whether or not they should accept the gamble (*“guesses”*). We also varied the outcome values and probability of the explicit gambles so that some had clearly positive or negative expected value (low uncertainty; *“easy decisions”*), while others had expected values close to ±0 (high uncertainty; *“hard decisions”*). This approach differs from many previous voluntary action studies, where participants are merely asked to make an action at a time they themselves choose (Kornhuber & Deecke, 1965; Libet, 1985; but see Khalighinejad et al., 2018). By embedding self-initiated actions in a classical economic decision-making paradigm, we provide a bridge between the study of voluntary action and that of decision-making and action more broadly.

Easy and difficult decisions here correspond respectively to “choosing” (deciding by considering reasons and evidence) to “picking” (deciding between two identical options, without reference to reasons or evidence) respectively (Ullmann-Margalit & Morgenbesser, 1977). Our *guess* trials, similar to our hard trials, are also a case of picking. However, decisions in the guess condition are based on an explicit instruction to generate action/non-action decisions internally, rather than equivocal external stimuli.

Our three conditions – easy decisions, difficult decisions, and guesses, can thus be organised along two independent dimensions (Table 1). First, they differ in the source, exogenous or endogenous, of the evidence that goes into the decision. If the RP reflects the activation of an endogenous route to action (Passingham, 1993), we should see a stronger RP signal on guessing trials than for either easy or difficult decisions. Second, our conditions differ in the strength of evidence, and thus the certainty of the decision. If the RP reflects decision uncertainty, we should see a stronger RP for the difficult decisions (expected value close to 0) than the easy decisions (expected value far from 0).

**Table 1.** Different types of action classified in terms of evidence source, and uncertainty.

| Evidence source | Uncertainty |  |
| --- | --- | --- |
|  | Low | High |
| Exogenous | Easy<br>decision | Difficult<br>decisions |
| Endogenous | - | Guesses |

## Methods

### Participants

Twenty participants (7 males, mean age = 25.5, SD = 4.7) completed the experiment, and were compensated £7.50 per hour for participation. The experiment lasted approximately 40 minutes. All procedures were approved by the UCL ICN ethics committee. This sample size was selected to match previous RP studies (Khalighinejad, Brann, Dorgham, & Haggard, 2019; Khalighinejad et al., 2018).

### Stimuli and Procedure

Participants were presented with 300 gambles, comprised of easy decisions, difficult decisions, and guesses. On the 212 decision trials, the probability of winning or losing, P(Win) and P(Loss), and the amount (value) that could be won or lost, V(Win) and V(Loss), were represented by a roulette-style wheel (Figure 1). P(Win) and P(Loss) were indicated by the size of the segments filled in green or red, respectively, while the amount that could be won or lost was shown numerically in each segment. The probability of winning varied from 20% to 80%, in 10% increments. The winning amount varied from +2 to +18 points in 2 point increments, while the losing amount was fixed at −10 points. The full set of probability/value combinations are shown in Supplementary Table S1, along with the expected value of each, calculated as P(Win) × V(Win) – P(Loss) × V(Loss). The average expected value across all bets was ±0. Gambles with an expected value of close to 0 were presented more often than those with extreme high or low values (see Table S1).

**Figure 1.**
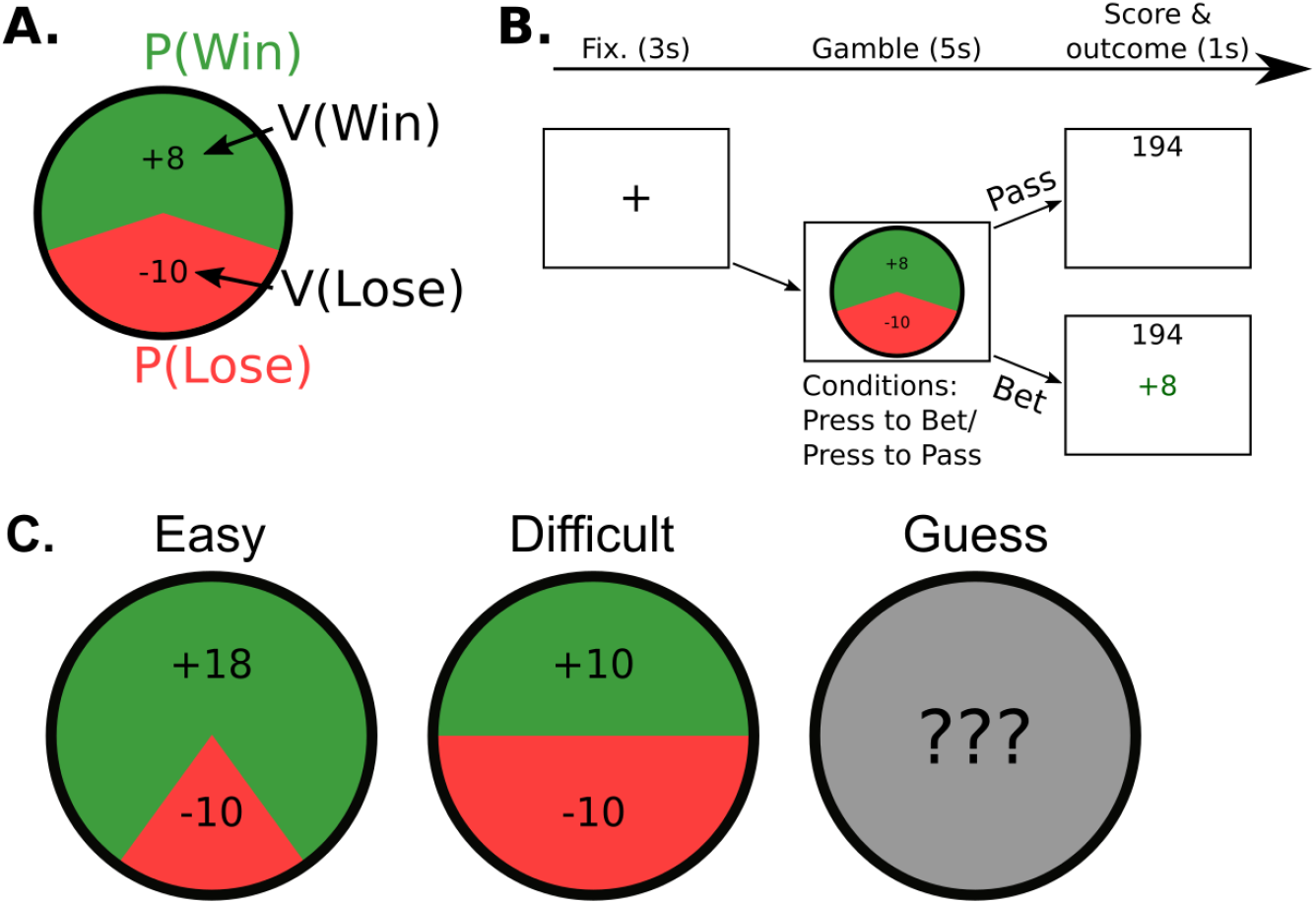
Stimuli and procedure. **A.** Gambles were presented visually in the form of a roulette wheel. The green and red areas indicate the probability of winning and losing respectively. Digits indicate the points that could be won or lost for the corresponding outcome. **B.** Each gamble was presented for 5 s. Participants either pressed the spacebar at any point during this window to accept the gamble, or to reject it (depending on the response condition). **C.** Examples of three types of gamble. Easy decisions had clearly positive or negative expected values. Difficult decisions had expected values close to 0, or at 0 in the case shown. For Guesses, the expected value on each particular trial was unknown.

Eighty-eight gambles out of the 300 were designated as guess trials. The gamble wheel on these trials was replaced by a grey circle, with text reading “???” inside (Figure 1). Participants were told that on these trials they would not see what the gamble was. Instead, they were told “On average, half of the guess gambles will be good, and half will be bad, so it’s totally up to you whether or not you want to take the risk on each one”, and “In the long run, you’re likely to break even on the guess gambles”.

After viewing each gamble, participants indicated whether they wanted to accept or reject it. In an *act-to-bet* condition, they pressed the spacebar within 5 seconds to accept the gamble, or withheld action to reject it – thus, rejection was the default. In an *act-to-pass* condition, the pressed the spacebar to reject the gamble and withheld action to accept it – acceptance was the default. By crossing these two response conditions with our other design factors, we can dissociate neural signals associated with positive or negative expected value from those associated with acting or withholding action (Guitart-Masip et al., 2014). After accepted gambles, participants saw outcome feedback showing the number of points that had been won or lost, in green or red font respectively, along with their cumulative points total, for 1 s. After rejected gambles, participants saw only their cumulative total for 1 s before proceeding to the next gamble.

Participants completed two practice blocks of 10 gambles each, one in each condition. In the experiment itself, participants completed four blocks of 75 gambles each. Response conditions alternated between blocks, and the first block (act-to-bet or act-to-pass was counterbalanced). For each participant, the same set of gambles were presented in blocks 1 and 2 and, again in blocks 3 and 4, in random order.

### EEG Acquisition

EEG was recorded using 32 + 6 channel BioSemi ActiveTwo system. We placed two reference electrodes on the mastiods, and monitored eye movements and blinks using electrodes placed on the outer canthi of each eye and above and below the right eye. During recording, EEG data were referenced to CMS-DRL, and offsets were maintained <30 μV. Data were recorded at 1024 Hz.

### Behavioural Analyses

All behavioural analyses were conducted in R. Mixed effects models fit using lme4 package (Bates et al., 2014). We fit a logistic mixed model for the probability of a bet being accepted on decision trials, ***logit(P(Bet)) ~ b_0_ + b_1_ P(Win) + b_2_ V(Win) + b_3_ V(Win) × P(Win) + b_4_ Condition***, where ***Condition*** is +1 for act-to-bet, −1 for act-to-pass. All parameters varied between participants as random effects. We tested a number of alternative models that performed worse in model comparison (see Results). Full details of all models can be found in Supplementary Materials, and R code can be found in the OSF repository accompanying the manuscript.

For each participant, we also sought to identify the decision trials that were sometimes accepted, and sometimes rejected (Figure 3B). We fit the same logistic regression models to each participant’s responses individually to estimate the probability, P(Bet) of each gamble being accepted by that participant, and transformed this quantity to obtain the probability of the participant acting on each particular trial, P(Act) = P(Bet) in the act-to-bet condition, and 1 – P(Bet) in the act-to-pass condition. We use this variable as a predictor in our EEG analyses, below. We identified the 50% of gambles closest to the point of indifference, where P(Bet) = .5, for each participant, and coded these as difficult decisions for that participant. We coded the remainder as easy decisions. This method takes into account how each participant weighted P(Win) and V(Win) to identify the trials that were most uncertain for that participant.

### EEG Preprocessing

Data were processed and analysed offline using custom python scripts and the MNE package for python (Gramfort et al., 2013). All channels were re-referenced to the average of the two mastoids. EEG segments with clear movement artefacts were removed prior to processing. We applied high pass (cut-off 0.05 Hz, width 0.1Hz) and low pass (cut-off 50 Hz, width 12.5 Hz) FIR filters, and a notch filter at 50 Hz to the raw data, and resampled to 125 Hz. ICA was used to identify and remove artefacts due to eye movements and blinks (Makeig et al., 1996).

On 96.4% of trials where an action occurred, it took place within 3 s of stimulus onset. Trials with slower response times than this were excluded from the analysis. Stimulus-locked epochs were extracted from -.5 s to +2 s relative to the onset of the gamble stimulus, and baselined between −0.1 and 0 s. Response-locked epochs were extracted from −2 s to +.5 s relative to the response time on trials where participants pressed the response button, and baselined between −2.1 s and −2 s prior to action. We excluded all epochs where voltages from any electrode exceeded ±120 μV from baseline. 7.7% of stimulus-locked epochs and 5.0% of response-locked were excluded in total.

In a typical RP experiment, actions are totally self-paced, and participants are not presented with any stimuli in the period prior to action. In this experiment, all of the actions analysed took place within 3 s of the onset of the gambles. This means that the 2 s window prior to action is often contaminated by stimulus-evoked activity. To isolate EEG components corresponding to motor preparation and execution from these stimulus-evoked components, we followed the procedure proposed by Kayser and Tenke (2006; Figure 2). Python scripts for this procedure are provided in the OSF repository accompanying the manuscript, and step-by-step details and visualisations are provided in Supplementary Materials.

**Figure 2.**
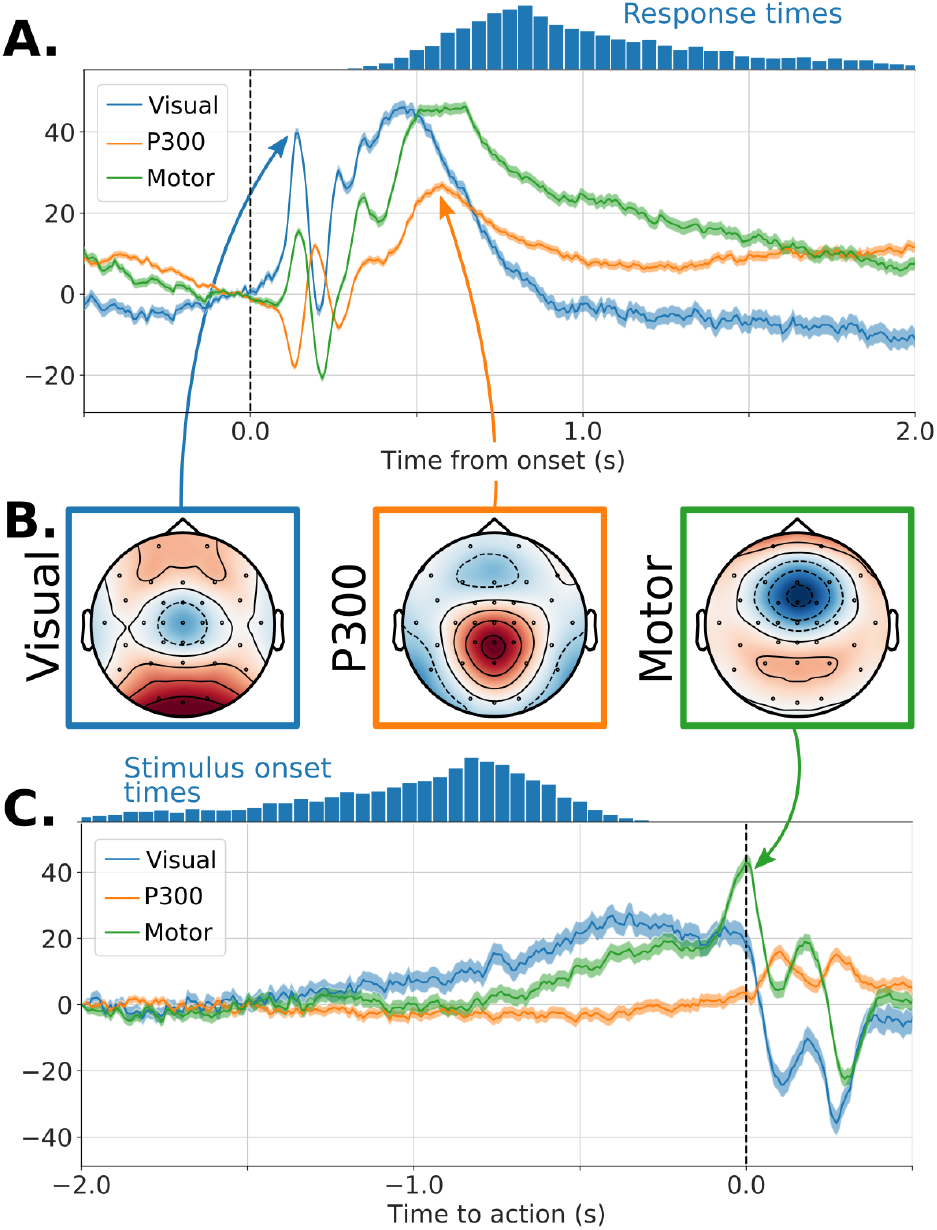
Time course and topography of the visual, P300, and motor components extracted by varimax PCA. **A.** Time course locked to stimulus onset. **B.** Spatial topography. **C.** Time course locked to the time of action.

We first estimated current source density (CSD) from the EEG using the surface Laplacian algorithm provided by Cohen (2014), with the Laplacian smoothing parameter to m = 5. This involves estimating the negative second spatial derivatives of the scalp EEG voltage, and effectively acts as a high-pass spatial filter, sharpening the topography of each component by reducing the blurring effects of volume conductance (Nunez et al., 2006). The raw EEG signal is dominated by the P300 component elicited by the onset of the stimulus, which is broadly distributed over centro-parietal electrodes (Supplementary Figure S4A-B). In the CSD signal, this component is restricted to a smaller region around CPz (Supplementary Figures S4C-D), revealing a negative prefrontal component around FCz that was obscured by the P300.

Next, we ran a Principal Components Analysis (PCA) on the CSD data, pooling across participants, using the data between −1 s and 0 s prior to action on trials where an action was produced. This decomposes the 32-channel EEG data into a collection of 32 orthogonal spatial components. Each component is defined by a eigenvector (weight vector), indicating how much activity at each electrode contributes to the component. In other words, the activity of each component is a weighted average of the activity recorded at each electrode.

PCA components are greedy: the first component captures as much variance as possible, the second captures as much of the variance not captured by the first component as possible, and so on. This means that each PCA component may not correspond to a single neural component, since the first component can capture variance that should be ascribed to later components. For this reason, we used varimax rotation (Kaiser, 1958). To do this, we select a subset of components with normalised eigenvalues great than 1 (Figure S5B) – that is, components that explain more variance than average. In this case, there were 9 such components, but consistent results were obtained using as few as 3 components. Varimax rotation finds the component weights that maximise the sum of squared loadings onto each component. This effectively redistributes the component weights so that variance is spread more evenly across components (Kaiser, 1958). The topographical distributions and time courses of the PCA and varimax components can be found in Figures S6 and S7. Note that varimax rotation is also commonly used in Factor Analysis, which estimates the weighting of individual survey items onto latent psychological components.

Finally, we inspected the rotated components to identify those that correspond to the neural components of interest, that we predicted might be present based on our task design. The three components shown in Figure 2 satisfied both a criterion of importance (i.e., variance explained) and interpretatibility (i.e., spatiotemporal pattern consistent with established neurophysiological studies, and with our task design). These were respectively interpreted as early visual processing, a centro-parietal positivity response (CPP, also know as the P300, see O’Connell, Dockree, & Kelly, 2012), and a negative motor-related component centred around electrode FCz. These components explained 18%, 14%, and 14% respectively of the variance in the response-locked EEG between −2 s and 0 s prior to action. In our analysis we focus on the motor preparation component, which we take to be analogous to the RP recorded prior to purely self-initiated actions.

### EEG Analysis

We noted that the shape of the response-locked motor preparation component depends on the response time on each trial, despite the steps taken to eliminate stimulus-evoked activity. This is caused by the initial stimulus-evoked peak in this component – a negative peak around FCz – approximately 500-600 ms after stimulus onset (Figure 2A). This is shortly after the peak of the P300 response. This peak is present even on trials where no action was executed, and occurs well before the average RT on trials where action did take place (see Figure S4), indicating that this activity is not due to actual motor execution. To explore the influence of this activity on our response-locked analyses, we plotted the motor component, locked to the time of action, separately across quartiles of the RT distribution (Figure 4A). The signal over the 100 ms prior to action is consistent across RT quartiles. Prior to this time, the shape of the motor preparation component depends on RT. On trials with rapid responses, the motor component ramps up from stimulus onset to the time of action. On trials with slower responses, early motor component activity peaks and begins to decline before the moment-related activity begins.

Based on this strong effect of response speed on the EEG ERPs, we included RT as a covariate in analyses of the motor component. To do this, we modelled the activation of the motor component over time in the window from 2 s to 0.1 s prior to action using a hierarchical generalised additive model, using lme4. We used a cubic splines basis matrix to capture the non-linear shape of the ERP over time (Hastie, Tibshirani, & Friedman, 2001). Based on BIC model comparison, we used regression splines with 5 knots. Interactions between all predictors and the basis matrix were included to capture the change in the ERP over time due to each predictor. For each predictor, the null hypothesis is that the shape of the ERP over time is unaffected by changes in the predictor. We test these null hypotheses using type II Wald χ^2^ tests, comparing the fit of models with and without these time × predictor interactions. This test is analogous to classical ANOVA. The code for this analysis can be found in the OSF repository accompanying the manuscript.

To test whether the motor preparation component is greater for difficult (uncertain) decisions than easy (certain) decisions, we fit a model to actions on decision trials. As a measure of certainty, we included P(Act), from close to 0.5 (high uncertainty) to 1.0 (low uncertainty) as a predictor. We also included response speed (inverse response times) and trial number as covariates. All predictors were z-transformed to obtain standardised regression weights. To test whether the motor preparation component is greater on guess trials than on difficult decisions, we fit a model to data from these two conditions, with a dummy coded predictor for condition, and the same covariates as above.

## Results

### Behavioural Results

On decision trials, participants were unsurprisingly more likely to accept gambles with higher P(Win), b = 4.03, SE = 0.31, z = 12.933, p < .001, and higher V(Win), b = 1.80 SE = 0.16, z = 11.400, p < .001 (Figure 3A). The influence of V(Win) was greater for higher values of P(Win), interaction b = 0.61 SE = 0.08, z = 7.630, p < .001. We found that participants’ responses were better fit by a four-parameter model, [P(Win), V(Win), their interaction, and an intercept term; AIC = 2797, BIC = 2860] than by two-(intercept and expected value; AIC = 3089, BIC = 3121) or one-parameter (expected value only; AIC = 3092, BIC = 3117) models. This indicates that participants relied on a heuristic weighted combination of each gambles attributes, and not the optimal decision-theoretic combination (see Supplementary Results for full model comparison; see also Rouault, Drugowitsch, & Koechlin, 2019). Participants were slightly more likely to gamble in the act-to-bet condition (51.8%) than the act-to-pass condition (47.4%), b = 0.50, SE = 0.10, z = 5.060, p < .001. Finally, in guessing trials, participants chose to bet on 49% of act-to-bet trials, SD = 30%, and on 46% of act-to-pass trials, SD = 26%, t(19) = 0.702, p = .491.

**Figure 3.**
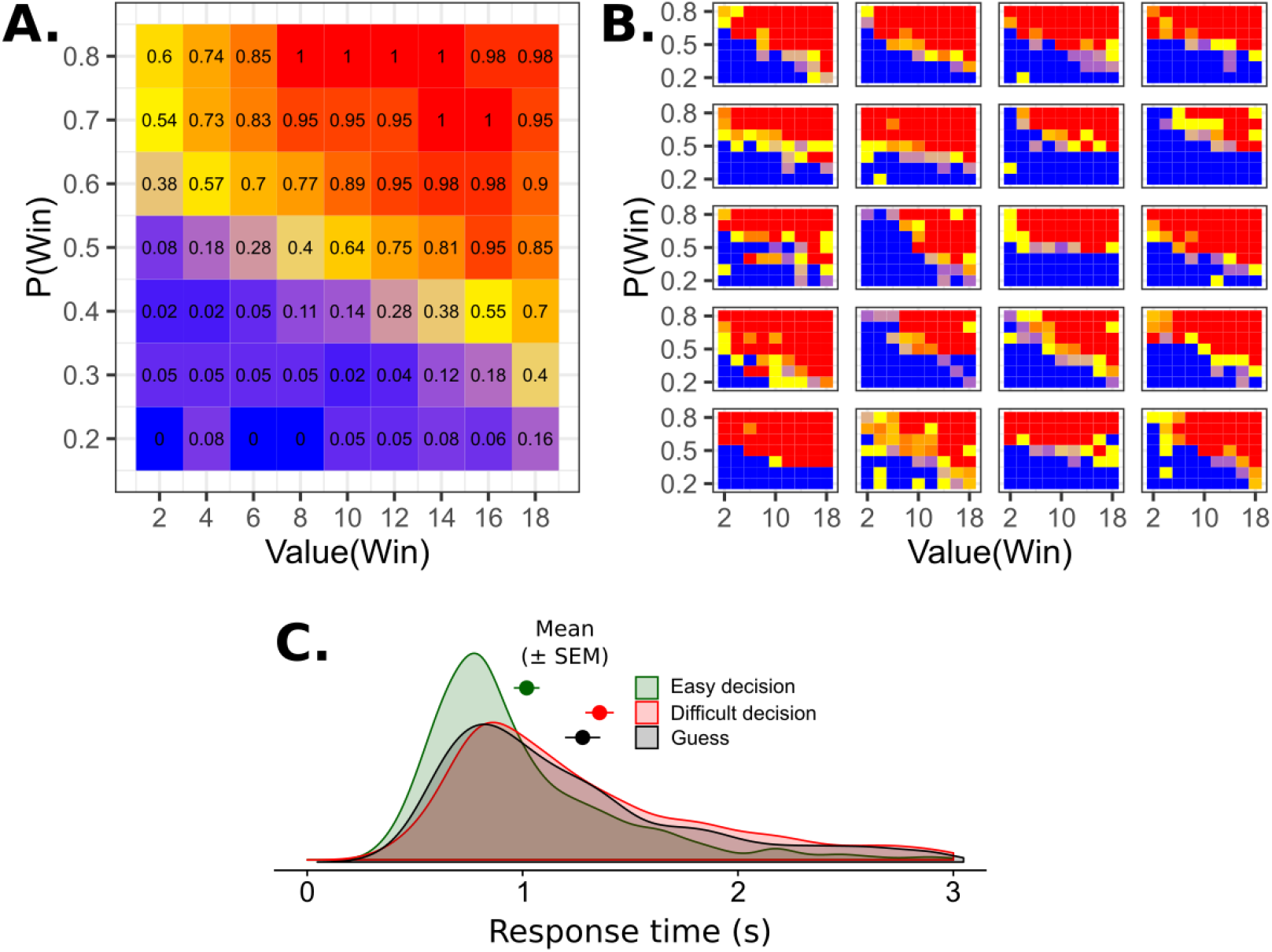
Participants’ decisions and response times. **A.** The probability of participants accepting each gamble, P(Bet), as a function of the number of points that could be won, Value(Win), and the probability of winning, P(Win). **B.** Choices for individual participants. Each participant was influenced by both Value(Win) and P(Win). **C.** Response times for easy decisions, difficult decisions, and guesses. Participants were significantly faster to produce actions for easy decisions than difficult decisions or guesses. Response times for difficult decisions and guesses did not differ.

Response times (RTs) followed a shifted log-normal distribution, consistent with an accumulation-to-threshold decision process (Figure 3C). There were very few RTs close to the 5 s limit, indicating that participants decided relatively early after stimulus presentation whether or not to act. Response times for certain decisions (1.02 s, SD = 0.27 s) were significantly faster than those for uncertain decisions (1.35 s, SD = 0.30 s), t(19) = 8.991, p < .001, or guesses (1.28 s, SD = 0.37 s), t(19) = 4.307, p < .001. Importantly, response times for uncertain decisions and guesses did not differ significantly, t(19) = 1.418, p = .172.

### EEG Results

We first test whether the motor component differs between actions for easy and difficult decisions. Mean ERPs for this test are shown in Figure 4B, with trials split by decision difficulty for visualisation. We found statistically significant effects of response speed, χ^2^(5) = 188.8, p < .001, and trial number, χ^2^(5) = 18.2, p = .003. However, we found no significant effect of certainty, χ^2^(5) = 6.5, p = .263. There was a significant certainty × response speed interaction, χ^2^(5) = 42.6, p < .001. This reflected the slightly higher activation for difficult decisions approximately 1.4 s prior to action on trials with slower RTs. This effect is only transient, and is eliminated by −1 s prior to action, indicating that it is unrelated to motor preparation. Therefore, we conclude that uncertainty does appreciably not affect the shape of the motor preparation component once differences in RT between trials have been controlled for.

**Figure 4.**
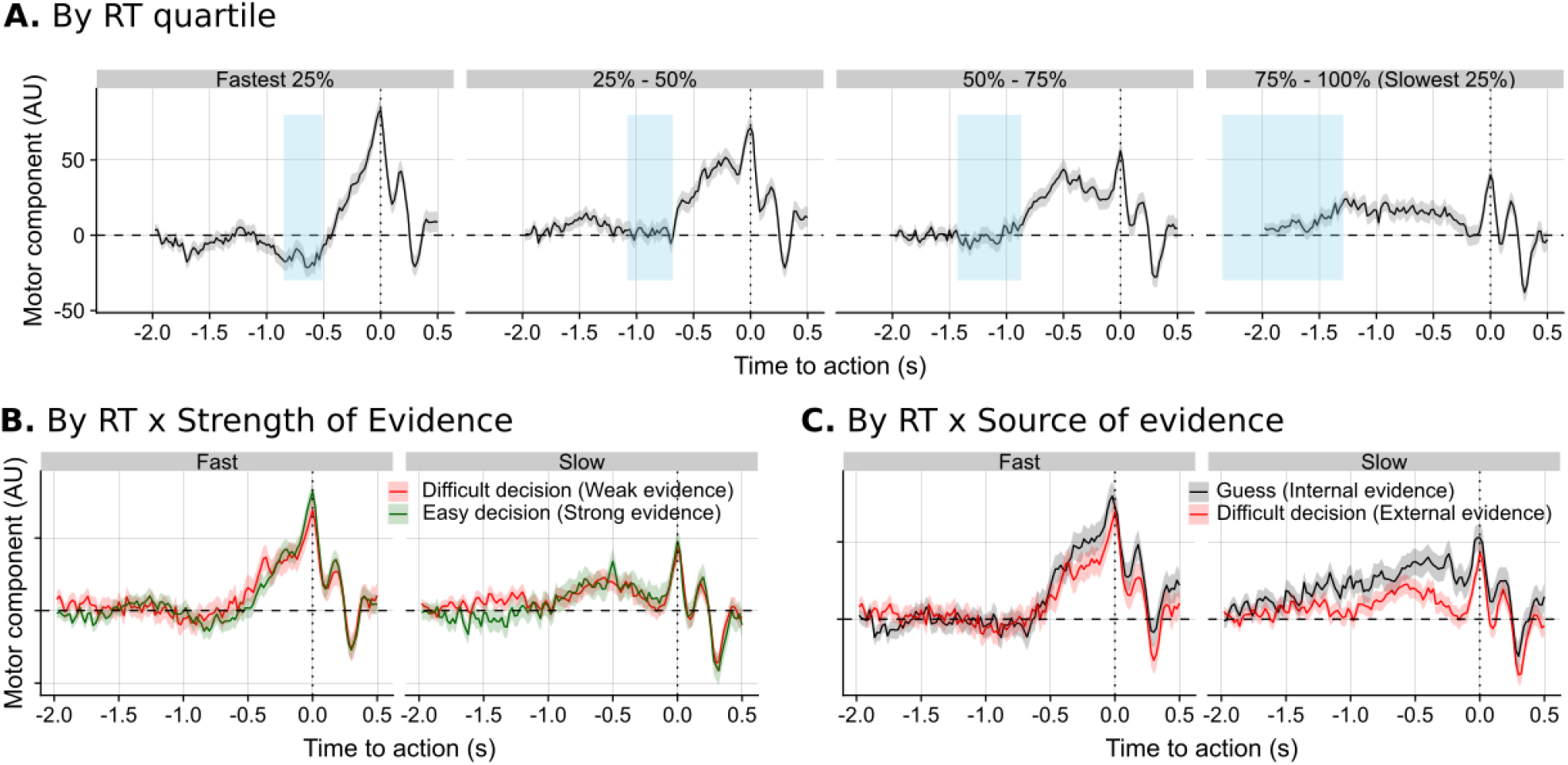
Action-locked event-related potentials. **A.** The shape of the action-locked motor component varied with response times. In trials with rapid responses, the ERP overlaps with stimulus-evoked activity, and produces a sharper ramp prior to action. Blue regions show mean ± 1 SD stimulus onset times within each quartile. **B.** Decision evidence strength did not affect the shape of the action-locked ERP after controlling for differences in response speed. **C.** Endogenously-produced actions on guess trials produced greater RP-like motor potentials than exogenously-produced actions on hard decision trials. This was particularly the case for slower responses.

We next tested whether the motor preparation component is greater for actions based on endogenous evidence (guesses) than actions based on uncertain exogenous evidence (difficult decisions). Recall that in both cases, participants’ sometimes decided to act, and sometimes to withhold action. In addition, response times did not differ significantly between the two conditions. However, participants must consider external evidence for hard decisions, but not when guessing. If the RP reflects decisions that are due to endogenous rather than exogenous evidence, we would expect to see a greater motor component activity in guessing than for difficult decisions. Mean ERPs for this test are shown in Figure 4C. We again found a statistically significant effect of response speed, χ^2^(5) = 161.40, p < .001. The effect of trial number was not significant, χ^2^(5) = 5.26, p = .385. Crucially, we also found a significant main effect of context, χ^2^(5) = 21.13, p < .001. In line with our predictions, the amplitude of motor preparation component was greater for guesses than for uncertain decisions. There was also a context × response speed interaction, χ^2^(6) = 41.2, p < .001. This reflects that the difference between contexts can be seen throughout the ERP on trials with slow RTs, but only at times closer to the action for trials with faster RTs (e.g. only after the onset of the stimuli). Therefore, we conclude that the shape of the motor preparation component is affected by the source of decision evidence, even when controlling for RT.

## Discussion

We sought to test whether the RP typically seen prior to voluntary actions reflects the fact that these actions are endogenously generated (Passingham, 1993; Passingham et al., 2010) or the fact that they are driven by noisy decision processes with high uncertainty (Nachev et al, 2005; 2008). Behavioural paradigms used to study the RP typically have no external triggering stimuli, and instead rely on a contextual instruction to “move when you feel like it”, or a purely internally-generated decision to move now (Khalighinejad et al., 2018). Experimental control of the conditions under which voluntary action decisions are made is therefore difficult. Here we use a novel combination of external stimuli from the established value-based decision-making field, and EEG filtering to extract RP-related signals under well-controlled conditions design to test contrast theoretical explanations of the causes of RP. Using a go/no-go gambling task and EEG filtering, we were able to isolate a negative-going motor preparation component that resembled the classical RP. We found that this component was more prominent prior to internally-generated actions than prior to actions driven by external evidence. This is consistent with the proposal that the RP reflects internal generation of action. We found that this RP-like component was largely insensitive to the level of evidence, and thus the degree of decision certainty. Specifically, we found no difference between actions driven by strong and weak external evidence, once differences in RT were controlled for. While caution is required in interpreting any null result, this finding appears inconsistent with theories that link the RP to uncertainty or conflict in decision-making.

The RP is often thought to be specific to ‘voluntary’ actions. However, it is often unclear just what is meant by a ‘voluntary action’. Fried, Haggard, He, and Schurger (2017) set out a number of key features of volition, which are useful in making sense of our results. In motor neurophysiology, voluntary actions have been defined as actions that occur without an external trigger (Passingham et al., 2010). In our task, every trial includes some external stimulus as participants must respond to the gamble presented. However, the decision-relevant information carried by the stimulus was systematically manipulated across trials. On standard trials, the stimulus both indicates that a decision must be taken (act or don’t act, bet or don’t bet) and provides the information to drive that decision: P(Win) and V(Win). On guess trials, the stimulus only indicates a decision must be taken. The information that drives the decision must be generated internally, for example by a spontaneous action decision, by an exploration strategy, or by retrieval from memory. We cannot identify exactly what processes underlay our participants’ guesses, but we argue that their actions on guess trials fulfil some of the criteria for voluntariness (Fried et al., 2017) more clearly than standard trials. Since the RP-like motor component occurred more clearly on guess trials than on trials with external evidence, we conclude that the RP is associated with internally triggered actions.

Some theories of volition emphasise the spontaneous or innovative character of voluntary actions: a voluntary action is often not predictable from the current context. This feature is consistent with the idea that voluntary actions are triggered by random fluctuations in the brain (Schurger et al., 2012). It is also consistent with the proposal that voluntary actions preferentially involve medial frontal cortex, because they involve cognitive conflict (Nachev et al., 2008). In our design, both guesses and difficult decisions were voluntary in this sense. In both cases, when presented with a single stimulus participants sometimes chose to bet, and sometimes chose to pass; they were somewhat random or unpredictable. Response times were also slower than easy decisions, indicating cognitive conflict. Following this logic, actions based on difficult decisions are more voluntary – less predictable, involve more conflict – than those based on easy decisions. This is consistent with another proposed feature of volition: an action is voluntary if the agent “could have done otherwise” (e.g. Hart, 1948). Importantly, however, we found that the more spontaneous actions in our difficult condition did not show a stronger RP-related motor component than the less spontaneous actions in our easy condition, once response times were controlled for. In contrast, we found strong differences between difficult decisions and guess trials, despite both of these classes of action being relatively spontaneous. On these grounds, we conclude that the RP is not a reliable marker of spontaneity or unpredictability of action decisions.

Our participants might have adopted suboptimal heuristics, for instance to bet on all trials where P(Win) > 0.5. There are a number of reasons why this seems unlikely. First, participants were slower to respond for difficult decisions than easy decisions. This would not be the case if participants followed a simple perceptual decision rule. Second, all participants responded inconsistently to at least some range of the probability/value space (shown as yellow regions in Figure 3B). Third, the responses of participants using a simple decision rule would be predicted by P(Win) or V(Win) only. We instead found that participants’ responses were influenced by P(Win), V(Win), and their interaction, and were broadly consistent with the expected value of the gambles shown. We conclude from these that participants made up their mind about each standard gamble on a trial-by-trial basis.

Similarly, participants might decide in advance whether to bet or not on the guess trials. We found that participants on average bet on roughly half of these trials, and 18/20 participants bet on between 15% and 85% of the 88 guess trials. The remaining four participants, who bet on 3, 4, 11, and 86 of the 88 guess trials, may have decided in advance how to respond when the guess prompt was presented. That is, they may have responded habitually to the guess trial stimulus, rather than making a de novo action decision on each guess trial. On that view, they might not be producing internally-generated actions on these trials. However, our results were unchanged when these participants were excluded. In fact, we would expect that including these participants should make us less likely to find a difference between guess trials and difficult decisions, so any predecision effects should count against the hypothesis linking RP to internal generation of action.

In this work, we report a number of methodological improvements which may be of use in future studies of voluntary action. To isolate EEG signals related to motor preparation from those elicited by the visual stimuli, we used a combination of surface Laplacian filtering and principle components analysis. While similar methods have been used in previous work (Kayser & Tenke, 2006)_-_to our knowledge this is the first application of these methods to extract RP-like signals from trials where sensory stimuli are also presented. Deconstruction of EEG to extract a response-related component when stimulus-related components are also present is consistent with the recent finding that preparatory signals are essentially similar for internally-generated and externally-triggered situations, though perhaps differing in accumulation rate (Elsayed et al, 2016; Lara et al, 2018). Further details are provided in the Supplementary Materials, and in the analysis code provided with this manuscript.

To model the shape of the motor component over time, we used hierarchical generalised additive models. These models are widely used to analyse non-linear time series in other domains, particularly in ecology (Pedersen, Miller, Simpson, & Ross, 2019), but to date have rarely been applied to neural time series. This technique is particularly useful for modelling slow, smooth components such as the RP and contingent negative variation. It may be less suited to modelling fast, non-linear components such as N1-P2, where a large number of basis functions are needed to capture the shape of the waveform. We plan to expand on this method in future work.

These models allowed us to control for response times in our analyses of the motor component. Differences in action latencies are a crucial but often overlooked confound in RP studies. It is well documented that the amplitude of the RP increases as participants wait longer before acting (Khalighinejad, Brann, Dorgham, & Haggard, 2019; Schurger, 2018; Verleger, Haake, Baur, & Śmigasiewicz, 2016). As a result, differences in RP amplitude between conditions are not interpretable unless differences in action latencies are controlled for, statistically or experimentally. This is a particular issue when trials begin with the presentation of a visual stimulus, as in this study, and others (Maoz et al., 2019). Unless perceptual and motor EEG components can be perfectly dissociated, apparent differences in the shape of the motor component may be artefacts caused by the onset or offset of visual components.

Finally, there is an important distinction between the RP and the lateralised readiness potential (LRP; Haggard & Eimer, 1999). The LRP is considered to be generated by the primary motor cortex contralateral to the hand used to act. While the RP reflects readiness to act in general, the LRP reflects preparation of a specific action. In the current experiment, participants responded only with the right hand. Classical methods of separating the RP from the LRP are based on a double-subtraction method, requiring responses from both hands (Eimer, 1998). Since we could not use those methods here, the later stages of our motor preparation component might include some component of LRP (Shibasaki & Hallett, 2006). Future work might test how our findings generalise to cases where participants must decide between two alternative actions, or between inaction and two possible actions. Such a design would make help to further disentangle readiness to act, conflict between competing actions, and internal and external sources of information.

## Conflict of interest

The authors declare no competing financial interests.

## Acknowledgements

This research was supported by a Research Project Grant from The Leverhulme Trust to PH (RPG-2016-378).The code to run this experiment, the preprocessed data, and the analysis code for this manuscript can be downloaded from https://osf.io/m834c/.

## Notes

### Competing Interest Statement

The authors have declared no competing interest.

