## Supplementary Materials for "The Readiness Potential reflects the internal source of action, rather than decision uncertainty"

### Stimuli

Table S1. Gamble stimuli used, and number times each was presented.

| Choice Gambles | | | | | | Guess Gambles | | |
|---|---|---|---|---|---|---|---|---|
| P(Win) | V(Win) | N | P(Win) | V(Win) | N | P(Win) | V(Win) | N |
| 0.2 | 2 | 1 | 0.5 | 10 | 4 | 0.5 | 10 | 4 |
| 0.2 | 4 | 1 | 0.5 | 12 | 3 | 0.5 | 12 | 3 |
| 0.2 | 6 | 1 | 0.5 | 14 | 2 | 0.5 | 14 | 2 |
| 0.2 | 8 | 1 | 0.5 | 16 | 1 | 0.5 | 16 | 1 |
| 0.2 | 10 | 1 | 0.5 | 18 | 1 | 0.5 | 18 | 1 |
| 0.2 | 12 | 1 | 0.6 | 2 | 1 | 0.6 | 2 | 1 |
| 0.2 | 14 | 2 | 0.6 | 4 | 2 | 0.6 | 4 | 2 |
| 0.2 | 16 | 2 | 0.6 | 6 | 3 | 0.6 | 6 | 3 |
| 0.2 | 18 | 3 | 0.6 | 8 | 3 | 0.6 | 8 | 3 |
| 0.3 | 2 | 1 | 0.6 | 10 | 2 | 0.6 | 10 | 2 |
| 0.3 | 4 | 1 | 0.6 | 12 | 2 | 0.6 | 12 | 2 |
| 0.3 | 6 | 1 | 0.6 | 14 | 1 | 0.6 | 14 | 1 |
| 0.3 | 8 | 1 | 0.6 | 16 | 1 | 0.6 | 16 | 1 |
| 0.3 | 10 | 1 | 0.6 | 18 | 1 | 0.6 | 18 | 1 |
| 0.3 | 12 | 2 | 0.7 | 2 | 2 | 0.7 | 2 | 2 |
| 0.3 | 14 | 3 | 0.7 | 4 | 3 | 0.7 | 4 | 3 |
| 0.3 | 16 | 3 | 0.7 | 6 | 3 | 0.7 | 6 | 3 |
| 0.3 | 18 | 2 | 0.7 | 8 | 2 | 0.7 | 8 | 2 |
| 0.4 | 2 | 1 | 0.7 | 10 | 1 | 0.7 | 10 | 1 |
| 0.4 | 4 | 1 | 0.7 | 12 | 1 | 0.7 | 12 | 1 |
| 0.4 | 6 | 1 | 0.7 | 14 | 1 | 0.7 | 14 | 1 |
| 0.4 | 8 | 2 | 0.7 | 16 | 1 | 0.7 | 16 | 1 |
| 0.4 | 10 | 2 | 0.7 | 18 | 1 | 0.7 | 18 | 1 |
| 0.4 | 12 | 3 | 0.8 | 2 | 3 | 0.8 | 2 | 3 |
| 0.4 | 14 | 3 | 0.8 | 4 | 2 | 0.8 | 4 | 2 |
| 0.4 | 16 | 2 | 0.8 | 6 | 2 | 0.8 | 6 | 2 |
| 0.4 | 18 | 1 | 0.8 | 8 | 1 | 0.8 | 8 | 1 |
| 0.5 | 2 | 1 | 0.8 | 10 | 1 | 0.8 | 10 | 1 |
| 0.5 | 4 | 1 | 0.8 | 12 | 1 | 0.8 | 12 | 1 |
| 0.5 | 6 | 2 | 0.8 | 14 | 1 | 0.8 | 14 | 1 |
| 0.5 | 8 | 3 | 0.8 | 16 | 1 | 0.8 | 16 | 1 |
| | | | 0.8 | 18 | 1 | 0.8 | 18 | 1 |

### Behavioural Analyses

To explore the factors that drove participants' decisions, we fit a number of logistic regression models to the responses given on non-guess trials (that is, trials where P(Win) and V(Win) were known). The models considered were as follows:

Table S2: Behavioural models fit.

| Model name | Link: logit[Pr(Bet)] = | N pars |
|---|---|---|
| Intercept only | $\beta_0$ | 1 |
| Amount only | $\beta_0 + \beta_1 \times V(Win)$ | 2 |
| Probability only | $\beta_0 + \beta_1 \times P(Win)$ | 2 |
| Expected value | $\beta_0 + \beta_1 \times [P(Win) \times V(Win) - P(Lose) \times V(Lose)]$ | 2 |
| Amount + Probability | $\beta_0 + \beta_1 \times V(Win) + \beta_2 \times P(Win)$ | 3 |
| Amount × Probability | $\beta_0 + \beta_1 \times V(Win) + \beta_2 \times P(Win) + \beta_3 \times V(Win) \times P(Win)$ | 4 |

For ease of interpretation, we centred P(Win) on it's mean value of 0.5 and V(Win) on it's mean of +10. In each case, the intercept parameter $\beta_0$ captures the bias towards betting ($\beta_0 > 0$) or passing ($\beta_0 < 0$), while the additional parameters $\beta_i$ capture the influence of predictor *i.* Note that with P(Win) and V(Win) centred, the *Expected value* model can be rewritten as

$$\text{logit}[\Pr(Bet)] = \beta_0 + \beta_1 \times [0.5 \times V(Win) + 20 \times P(Win) + 1 \times V(Win) \times P(Win)]$$

This means that the *Expected value* model is a special case of the *Amount × Probability* model, with the ratio of the coefficients $\beta_1{:}\beta_2{:}\beta_3$ fixed at 0.5:20:1. Similarly, the *Amount + Probability* is a special case of the *Amount × Probability* model with the interaction coefficient $\beta_3$ set to 0.

We first fit each model to each participant individually. Figure S1 shows the observed probability of betting for each combination of P(Win) and V(Win) for each participant, along with the predictions of each of our models, fit to that participant's data. To evaluate model fit for each participant, we first calculated the Bayesian Information Criterion (BIC) = $\log(n)k - 2\log(L)$, where $n$ is the number of data points, $k$ the number of parameters, and $L$ is the likelihood of the model. Figure S2 shows the posterior model probability of each model, for each participant, calculated from BIC scores (Wagenmakers & Farrell, 2004). This shows that most participants' data are best captured by the *Amount × Probability*, with some participants better modelled by the *Amount + Probability* or *Expected value* models. For most participants, the data are well explained by two or three of these models, indicating that it is difficult to differentiate between these models at an individual level.

We next examined the parameters of the full *Amount × Probability* model fit for each participant (Figure S3). In order to compare this model to the restricted *Expected value* model, we first normalised the regression weights $\beta_1$, $\beta_2$, $\beta_3$ to sum to 21.5 for each participant. As noted above, if participants use the *Expected value* model, we would expect these normalised coefficients to have means of 0.5, 20, and 1 respectively. We found the average regression weights across participants did not differ significantly from these predictions for any of these predictors (Table S3). On average, the influence of P(Win) was greater than would be expected under the *Expected Value* model, while the influence of V(Win) and the interaction are less.

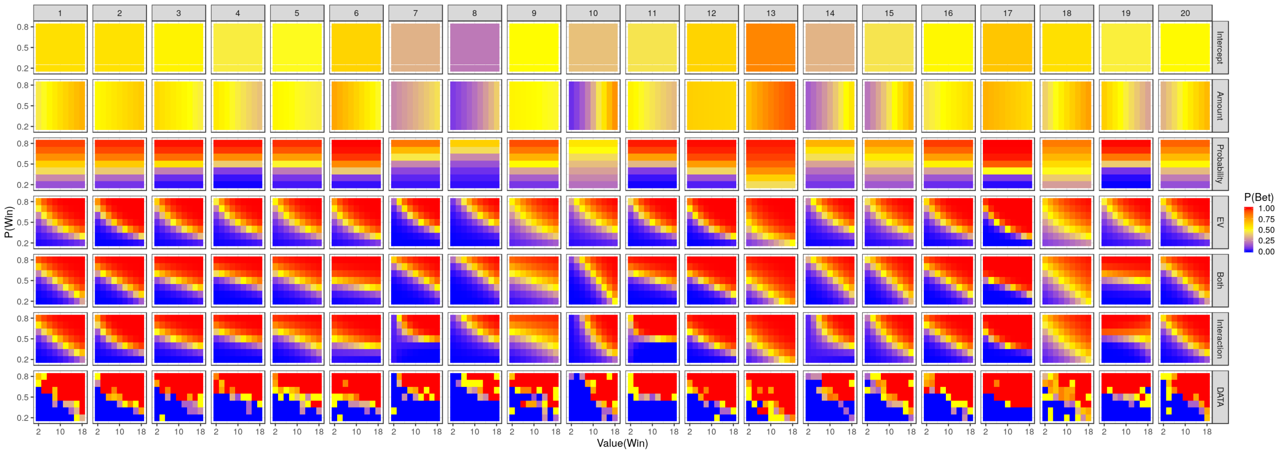


*Figure S1. Actual responses by each participant (bottom row), along with predicted responses for each model, fit to that participant's data.*

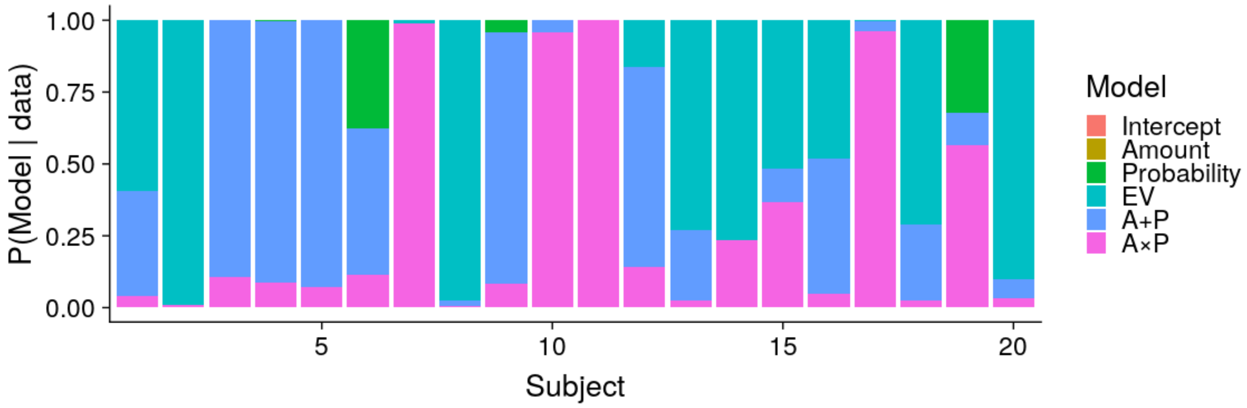


*Figure S2. Posterior model probabilities for each participant, estimated from BIC scores.*

Taken together, these results indicate that while participant's behaviour is broadly consistent with an *Expected Value* model on average, individual participants differ from this model in systematic ways. We confirmed this interpretation using hierarchical regression modelling, allowing parameters to vary between participants. We began by fitting the models above as hierarchical logistic regression models using the lme4 package for R, and all parameters were allowed to vary across participants. As noted in the main manuscript, model comparisons strongly supported the *Amount* × *Probability* model over all of the alternatives (Table S4).

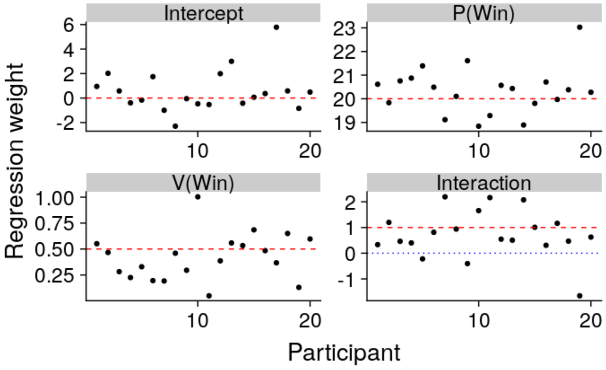


Figure S3. Estimated normalised regression weights for each participant were distributed close to the values that would be expected if decisions were driven by the Expected Value of the gambles (red lines).

Table S3. Individual participants' regression weights did not differ significantly from those predicted by Expected Value theory.

| Term | Mean | SD | SEM | Reference | t-value | p-value |
|---|---|---|---|---|---|---|
| Intercept | 0.57 | 1.73 | 0.39 | 0 | 1.469 | 0.158 |
| P(Win) | 20.35 | 0.98 | 0.22 | 20 | 1.584 | 0.130 |
| V(Win) | 0.42 | 0.25 | 0.05 | 0.5 | -1.546 | 0.139 |
| Interaction | 0.73 | 0.97 | 0.21 | 1 | -1.321 | 0.202 |

Table S4. Model comparison for the hierarchical regression models.

| Model | LL | DF | AIC | BIC |
|---|---|---|---|---|
| Intercept only | -2827.38 | 2 | 5658.76 | 5671.47 |
| Amount only | -2771.32 | 5 | 5552.64 | 5584.40 |
| Probability only | -1822.85 | 5 | 3655.70 | 3687.46 |
| Expected value | -1539.58 | 5 | 3089.16 | 3120.92 |
| Amount + Probability | -1418.57 | 9 | 2855.15 | 2912.32 |
| **Amount × Probability** | **-1388.58** | **10** | **2797.17** | **2860.69** |

### Isolating EEG Components

Traditionally, the Readiness Potential is recorded in the seconds prior to self-initiated actions. This means that no stimuli were presented immediately prior to the action in question, and as a result the EEG does not contain stimulus-evoked activity. In the current experiment, each trial begins with the a gamble being shown on screen, and responses typically occurred 1-2 s after the gamble was presented. As a result, the EEG prior to the action is contaminated by components evoked by the stimuli.

To isolate EEG components related to motor preparation, we adapt a procedure introduced by (Kayser & Tenke, 2006a, 2006b). We first apply a Surface Laplacian filter to the data to help localise EEG components. This transforms the data from a measure of electric potentials at the scalp to an estimate of underlying current sources and sinks: the Current Source Density (CSD). The Surface Laplacian can also be thought of as a spatial high-pass filter applied to the data, which attenuates low-spatial-frequency signals that are broadly distributed across the scalp, but preserves high-spatial-frequency signals that are more localised (Cohen, 2014). We used the surface Laplacian algorithm provided by Cohen (2014), reimplemented in python. The Laplacian smoothing parameter was set to m = 5. Figure S4 shows the original EEG signal (top) and current source density estimates after applying the Surface Laplacian (bottom) for stimulus- and response-locked epochs. The EEG signal is dominated by a strong P300 response, peaking approximately 500 ms after stimulus onset. This component is attenuated in the CSD data, revealing a negative component around electrode FCz, peaking just after the P300 in the stimulus-locked epochs, and at the time of action in the response-locked epochs.

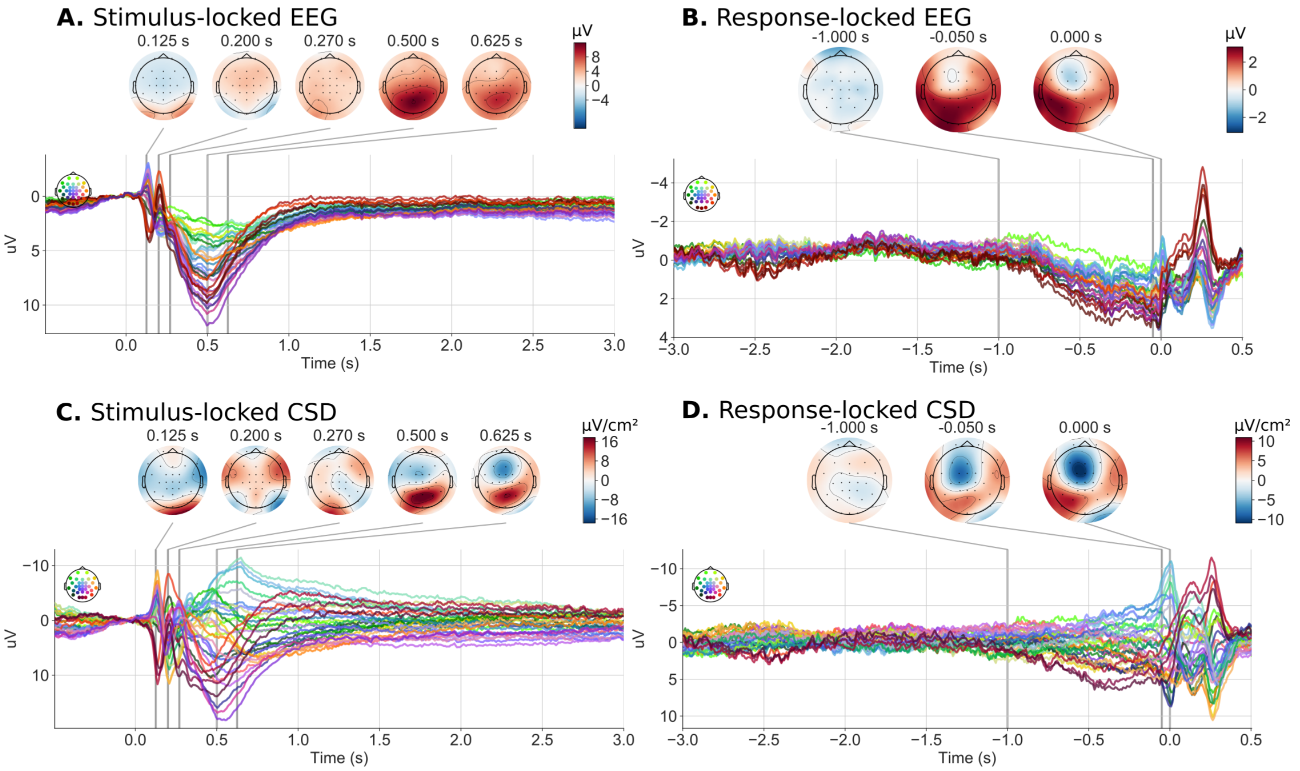


*Figure S4. The original EEG signal (top) and Current Source Density estimates (bottom) after application of the Surface Laplacian, for stimulus- (left) and response-locked (right) epochs.*

Next, we conducted Principle Components Analysis (PCA) on the covariance matrix for the CSD data between 1 second prior to action and the time of action itself (Figure S5A). This covariance matrix was calculated separately for each trial, and then averaged across trials. Nine of the thirty-two PCA components had standardised eigenvalues greater than 1 (Figure S5B & C). We retained these nine components for varimax rotation, resulting in the rotated components shown in Figure S5D. The stimulus- and response-locked time courses of the unrotated PCA components are shown in Figure S6, and those for the components after varimax rotation are shown in Figure S7. We used these time courses and scalp topographies to identify varimax component 2 as our putative motor preparation component, analogous to the Readiness Potential recorded prior to self-initiated actions.

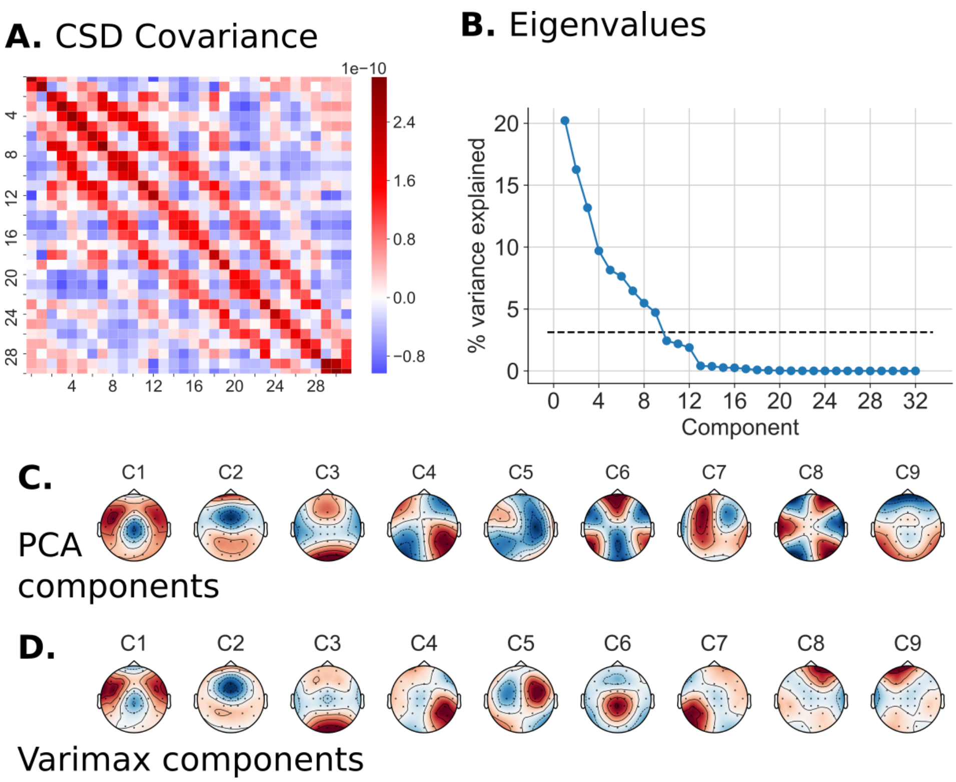


*Figure S5.* ***A.*** *Averaged covariance matrix used for PCA.* ***B.*** *% variance explained by each PCA component. Vertical line corresponds to a standardised eigenvalue of 1.* ***C.*** *Scalp topographies for the 9 PCA components with standardised eigenvalues > 1.* ***D.*** *Topographies for these components after varimax rotation.*

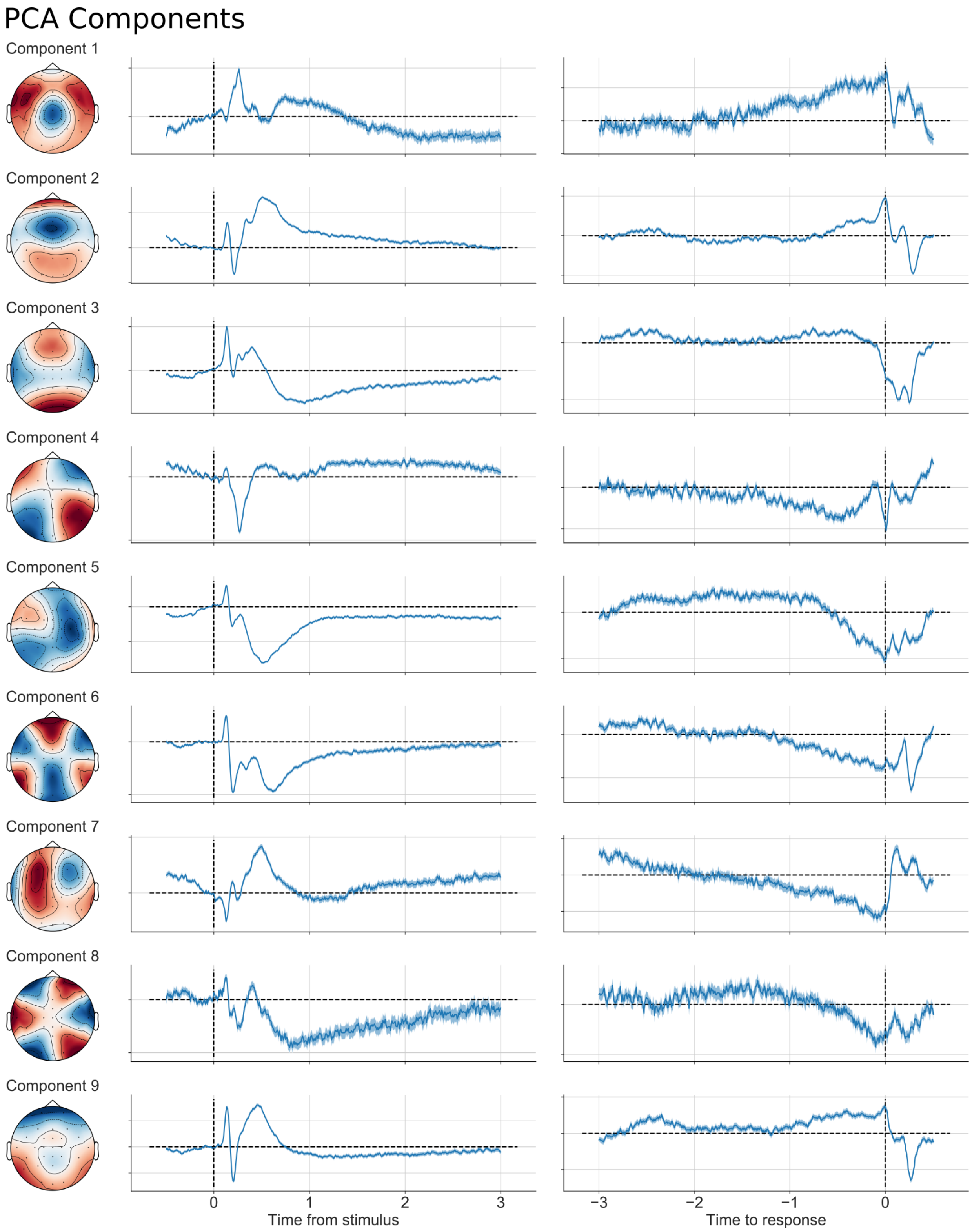


*Figure S6. Stimulus- and response-locked time courses of unrotated PCA components.*

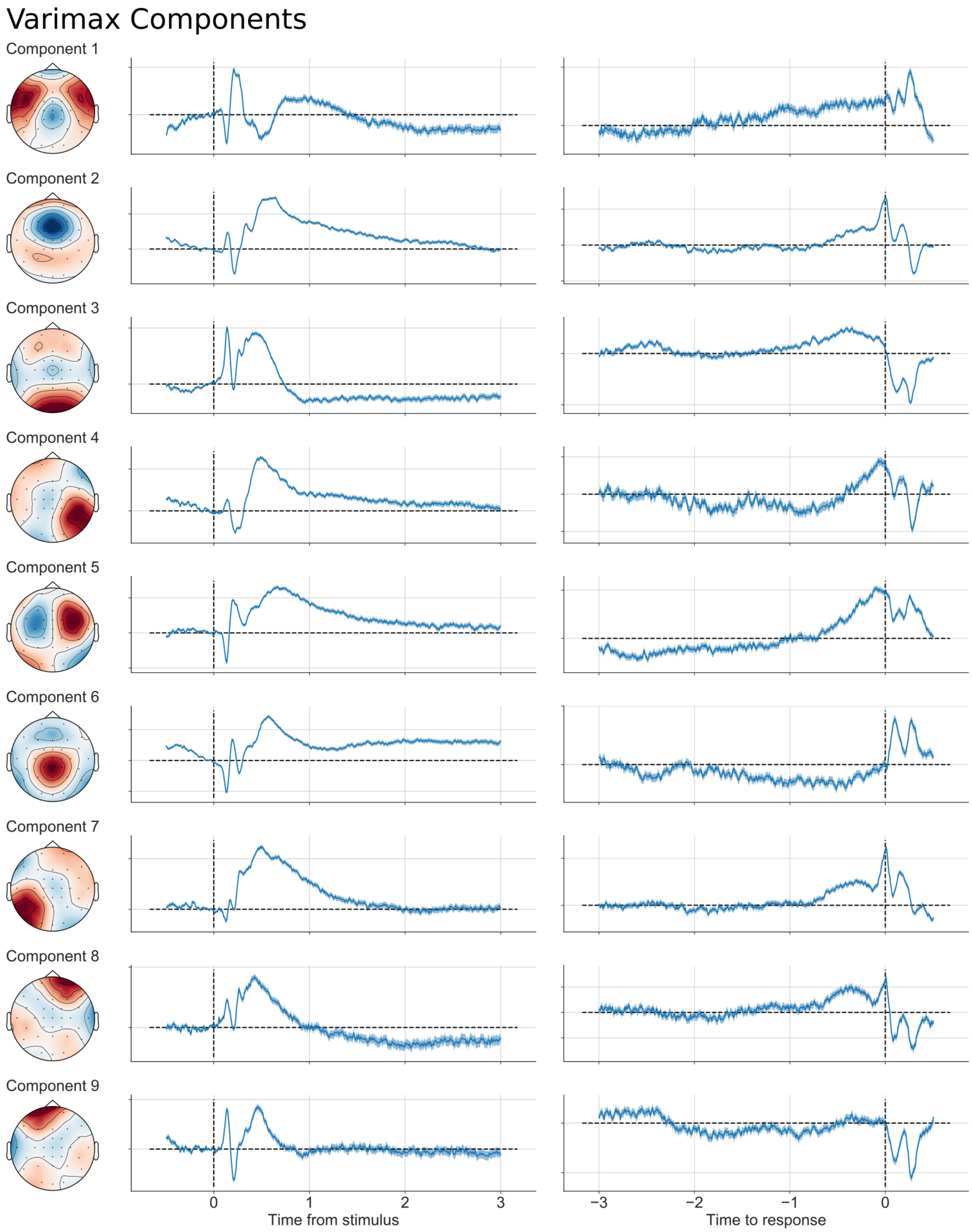


*Figure 7. Stimulus- and response-locked time courses of the components after varimax rotation. The primary analysis reported in the manuscript are conducted on Component 2.*
